# UCE phylogenomics inform the systematics and geographic range evolution of the harvester ant genus *Pogonomyrmex*

**DOI:** 10.1101/2024.11.12.623263

**Authors:** Leland C. Graber, Robert A. Johnson, Corrie S. Moreau

## Abstract

The harvester ant genus *Pogonomyrmex* is common in arid regions of North America, South America, and Hispaniola. Here, we use ultraconserved elements (UCEs) to infer the most comprehensive phylogeny of the genus to date. Our inferred topology largely supports previously hypothesized *Pogonomyrmex* evolutionary relationships and species groups. Our divergence dating analysis substantially pushes back the age of both *Pogonomyrmex* and the Pogonomyrmecini tribe, suggesting an early Eocene origin for *Pogonomyrmex*. We infer that *Pogonomyrmex* likely originated within the Pacific dominion ecoregion of Northern South America. *Pogonomyrmex* (excluding the *Pogonomyrmex mayri* clade) likely adapted to cooler and drier climates in the middle Eocene. The genus spread to North America in the late Eocene, likely aided by the ephemeral volcanic islands of the Panama Arc. A smaller group of *Pogonomyrmex* dispersed to Mesoamerica and North America in the Oligocene and to Hispaniola in the Miocene, most likely out of Mesoamerica. Our results suggest that adaptations to arid environments allowed *Pogonomyrmex* to spread and diversify in regions of the Americas that were actively becoming cooler and drier due to orogeny and other geologic processes.

## Introduction

*Pogonomyrmex* Mayr, 1868 is a genus of harvester ant common in arid habitats in the Americas (Johnson and Cover 2015). The genus name translates to “bearded ant,” a reference to the psammophore, the coarse hairs on the ventral surface of the ant’s head that are used to carry sand or soil in the excavation of the below ground nest. The genus currently has 95 valid species, 2 subspecies, and 1 fossil species from the Florrisant Formation in Colorado, USA (Johnson 2021). *Pogonomyrmex* is found in three non-overlapping faunal groups in North America, South America, and on the island of Hispaniola (Johnson and Cover 2015). *Pogonomyrmex* ants feed predominantly on seeds, though many species feed on dead arthropods as well (MacMahon, Mull, and Crist 2000)

The highest concentration of species diversity of *Pogonomyrmex* is in the Mojave, Sonoran and Chihuahuan deserts of the southwest United States and northwest Mexico, with many species co-occuring in the same areas (Guénard et al. 2017) (Figure 1). While not as concentrated, the deserts of northwest and west-central Argentina have nearly as much species diversity as the deserts of North America, though they are historically less studied than their North American counterparts. Recent work in southern Argentina and Chile has revealed many more South American species than previously known; two-thirds of described *Pogonomyrmex* species are found in South America (Johnson 2015, 2021). While *Pogonomyrmex* is most prevalent in deserts, species are found in the temperate grasslands of western North America as far north as southern Canada (Glasier et al. 2013). Only one species, the dimorphic *Pogonomyrmex badius*, is found on the coastal plains of the southeastern United States (Cole 1968). Two species, *Pogonomyrmex guatemaltecus* and *Pogonomyrmex humerotumidus*, are found in Mesoamerica, and three species are found on the island of Hispaniola (Vásquez-Bolaños and Mackay 2004). In South America, a small number of *Pogonomyrmex* species are found outside of deserts or arid grasslands. *Pogonomyrmex sylvestris*-group species inhabit high elevation cloud forests in Venezuela, Colombia, and Ecuador (Lattke 2006). *Pogonomyrmex mayri* has a patchy distribution within the dry forests and coastal xeric scrub of Colombia. *Pogonomyrmex naegelii* is the most cosmopolitan *Pogonomyrmex* species, occurring broadly across South America except in deserts, rainforests, and high-elevation areas (Johnson 2015).

**Figure 1:**
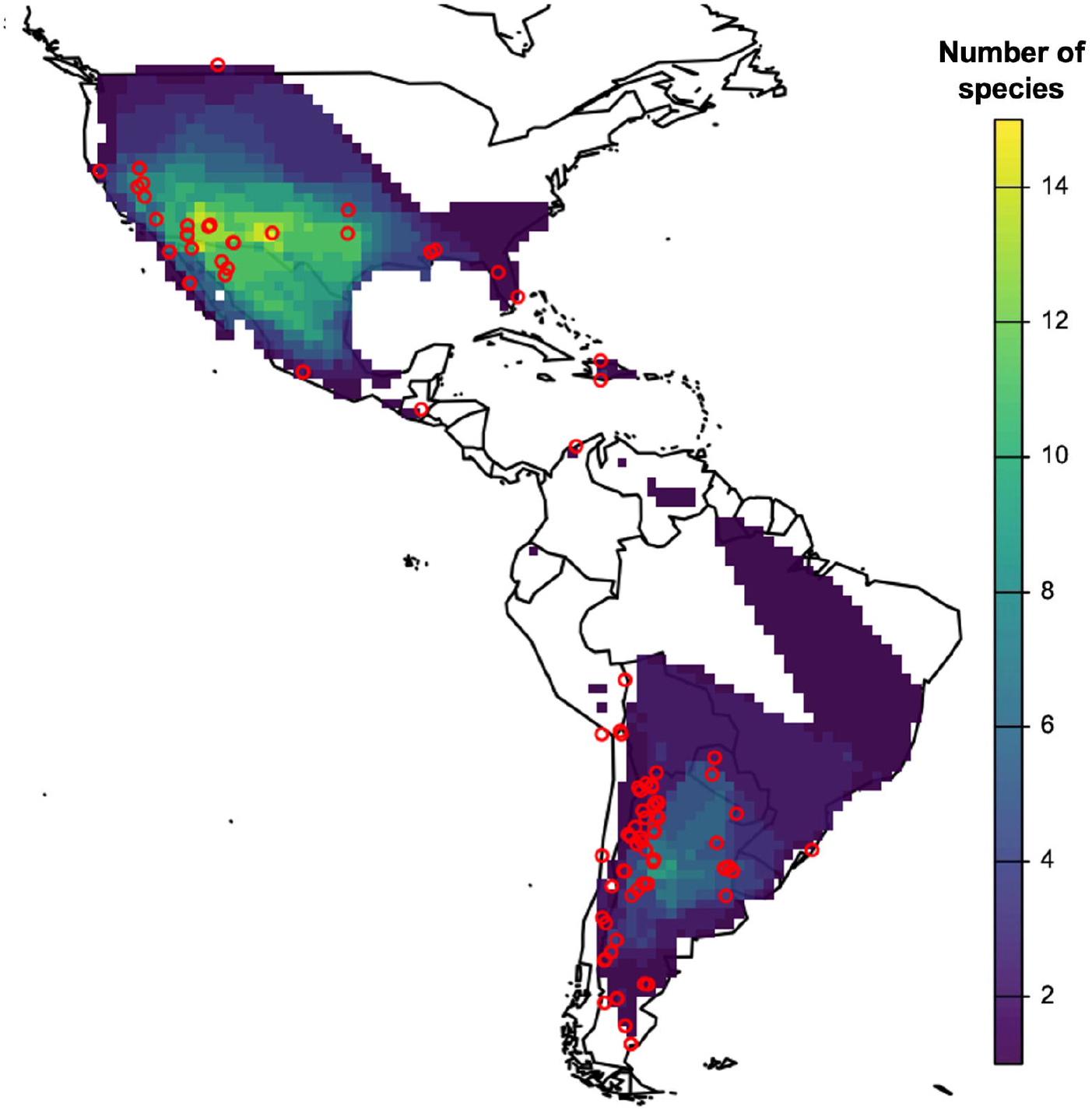
*Pogonomyrmex* species richness map. Individual species ranges for each of the 93 species were constructed using collection locality data from the Global Ant Biodiversity Informatics (GABI) database (Guénard et al. 2017) and research grade observations from iNaturalist (https://www.inaturalist.org). Predicted ranges were estimated for species without sufficient observational data using available occurrence data and area geography. Red dots indicate the collection locations of samples used in this study with exact latitude and longitude coordinate data. Colors correspond to species diversity across their range with cooler colors (purple) representing the least number of species and warmer colors (yellow) higher number of species in a given location.

While no formal historical biogeographic analysis has been done of the genus, competing explanations have been proposed to explain the geographic history of *Pogonomyrmex* and the tribe Pogonomyrmecini, which, in addition to *Pogonomyrmex*, contains *Hylomyrma* and *Patagonomyrmex*, a more recently described genus made up of species thought to be *Pogonomyrmex* until recently (Johnson and Moreau 2016). Wheeler (1914), and later Creighton (1952), hypothesized that the genus originated in the boreal forests of North America after splitting from the ancestors of the genus *Myrmica*. Kusnezov (1951) was the first to propose a South American origin for the group, arguing that it was unlikely for Nearctic *Myrmica* species and *Pogonomyrmex* (then including genera *Hylomyrma* and *Patagonomyrmex*) to have shared a common ancestor and that the divergence of the groups occurred much earlier than Wheeler (1914) and Creighton (1952) believed (Fernández and Palacio 1997).

The phylogeny of *Pogonomyrmex* has been inferred previously using both morphological characters and Sanger-sequenced genes, albeit with incomplete species sampling (Johnson and Moreau 2016; Parker and Rissing 2002; Taber 1999). To further investigate evolutionary relationships within *Pogonomyrmex*, as well as to formally infer the genus’s age and biogeographic history, we leveraged phylogenomic methods. Phylogenomic approaches incorporate many more phylogenetic markers than previous methods and are becoming increasingly more common in studies of insect systematics and biogeography, largely due to increased availability and affordability of high-throughput sequencing methods. Phylogenomic methods have been shown to resolve many previously unresolvable insect clades (Cruaud et al. 2021; Van Dam et al. 2017), and are increasingly being adopted in studies of ant evolution (Borowiec et al. 2024; Price et al. 2022; Ješovnik et al. 2017; Philip S Ward et al. 2015).Ultraconserved elements (UCEs) are one such phylogenomic approach commonly used in ant systematics; the most recent ant-specific UCE probe set targets 2,590 UCE loci (Branstetter et al. 2017). Additionally, it is possible to get usable UCE sequence data from less than optimally preserved material, such as pinned specimens (Blaimer et al. 2016).

In this study, we use ultraconserved elements (UCEs) to infer a new phylogeny for *Pogonomyrmex* to better understand the relationships between species in the genus as well as the relationships between *Pogonomyrmex* and the other genera in the tribe Pogonomyrmecini. We used the newly inferred phylogeny and a secondary calibration to infer the divergence dates of *Pogonomyrmex* evolution and then used the resulting dated tree and geographic range data to infer the geographic origins of *Pogonomyrmex* and evolution of the genus’s current geographic range.

## Materials and Methods

### Taxon sampling

Specimens included in this study were collected between 1924 and 2018 from locations across the geographic range of *Pogonomyrmex* (Figure 1). Most specimens collected were preserved in 95% ethanol, though a minority of specimens had been dried and pinned. The initial dataset included 286 *Pogonomyrmex* samples; however, due to contamination and poor sequence quality in many samples, the final dataset represented in the phylogeny includes 143 *Pogonomyrmex* samples and 11 outgroups. The dataset includes six not-yet-described *Pogonomyrmex* species (species that are not yet described are indicated by an “RAJ” followed by a short working name). Outgroup sequences were obtained from the SRA database at NCBI and include some species from Pogonomyrmecini tribe members *Hylomyrma* and *Patagonomyrmex*, as well as more distantly-related Myrmecines *Myrmica, Veromessor*, and *Aphaenogaster*.

To test for monophyly of species we included multiple individuals from species previously found to be cryptic or polyphyletic (Johnson and Moreau, unpublished). These multiple individuals for cryptic species were chosen to represent a variety of collection locations. Specimen codes, specimen age, preservation method, locality data and number of UCE loci recovered can be found in the Supplementary Tables.

### Library preparation, target enrichment, UCE sequencing

DNA extractions were conducted at the Field Museum’s Pritzker DNA laboratory from 2008 to 2011 and at Cornell University in 2020, 2021, and 2023. The majority of DNA was extracted destructively from whole ant specimens or legs preserved in ethanol using the DNEasy/MoBio PowerSoil Kit (Qiagen). A smaller number of DNA samples were extracted non-destructively using the DNEasy Blood and Tissue Kit (Qiagen) from pinned specimens—specimens were left intact, save for a small hole pierced in the thorax (P.S. Ward, 2019 protocol). A Qubit fluorometer (Life Technologies Inc.) was used for DNA quantification after extraction. Samples that were destructively extracted were diluted appropriately and sheared to a target size of 500 bp using a sonicator (Covaris). This sheared DNA, as well as non-sheared DNA from non-destructively extracted specimens, was used as input for library preparation.

Library preparation was performed at Cornell University in 2020, 2022, and 2023, and followed a modified genomic DNA library preparation protocol (Kapa Hyper Prep Library Kit, Kapa Biosystems, Faircloth et al. 2015) that used “speedbeads”, a generic SPRI bead (Rohland and Reich) and custom iTru dual-indexing barcodes (Glenn et al. 2016). After end repair and a-tailing of extracted sequences and a ligation reaction with annealed stubs, dual indices were added, and library amplification was performed. Libraries were then pooled (4-12 samples/pool) at equimolar ratios based on similarities of post amplification Qubit values. Pooled libraries were then enriched with a set of 9,446 probes (MYcoarray, Inc.) targeting 2,524 ant-specific UCE loci (Branstetter et al. 2017b). We adhered to the library enrichment procedures outlined in the Arbor Biosciences MYBaits kit (Blumenstiel et al. 2010), utilizing a 0.2X dilution of the standard probes. For PCR recovery of enriched libraries, we employed a with-bead method (Faircloth et al. 2015). Next, a purification step was performed using 1.0× speedbeads. The resulting enriched pools were reconstituted in 30 µl of elution buffer. Subsequently, these pools underwent analysis on an Agilent TapeStation 4150 Instrument to determine their concentration; all pools that displayed sufficient concentrations (at or above 4 nM) and were incorporated into the four final consolidated pools. Sequencing of the combined library pools occurred over six distinct 150-bp paired-end Illumina sequencing runs. The first two and last two runs took place on a NextSeq 500 instrument at the Cornell University Biotechnology Resource Center, and the middle two runs took place on a HiSeq 4000 instrument at Admera Health in South Plainfield, New Jersey. Data was demultiplexed by the Cornell University Biotechnology Resource Center and Admera Health after each provided sequencing service.

### Trimming, processing, and alignment of UCE data

All computation steps were performed on remotely accessed computers at the Cornell Institute of Technology’s BioHPC Cloud. For initial sequence processing, we used the Phyluce software, which allows for use of other packages within the Phyluce environment (Faircloth 2016). Illumiprocessor (Faircloth 2013), a parallel wrapper for the Trimmomatic package (Bolger, Lohse, and Usadel 2014) was used to trim the demultiplexed FASTQ data by removing low-quality bases and adaptor indexes. The assembly of cleaned reads was carried out through parallel wrappers around the SPAdes software (“phyluce_assembly_assemblo_spades”) (Bankevich et al. 2012). Contig assemblies were aligned with a FASTA file containing all ant-specific hymenoptera baits (“phyluce_assembly_match_contigs_to_probes”) which enabled the identification of contigs corresponding to enriched UCE loci for each taxon. Sequence coverage statistics for contigs containing UCE loci were also computed. FASTA files for individual loci were constructed that contained sequence data specific to the taxa present at each locus; the FASTA files were then aligned with MAFFT (“phyluce_align_seqcap_align”) (Katoh, Asimenos, and Toh 2009). Alignments were internally trimmed (“phyluce_align_get_gblocks_trimmed_alignments_from_untrimmed”) using a wrapper around GBlocks (Castresana 2000) and a relaxed setting, with b1-b4 parameters specified as 0.5, 0.5, 12, and 7, respectively.

### Contaminant detection and quality control of UCE data

Inspection of initial alignments and early tree inferences revealed obvious problems in aligned sequence data, suggesting sequence contamination or other quality issues. As a first contamination check, a Phyluce function (“phyluce_assembly_match_contigs_to_barcodes”) and reference barcodes for legacy genes *COI, wnt*, and *LwRh* obtained from GenBank (National Center for Biotechnology Information, National Library of Medicine) were used to identify homologous genes in our concatenated UCE sequences. The pulled-out genes were then put into BLAST (National Center for Biotechnology Information, National Library of Medicine) and compared with database sequences in order to identify contamination.

Since not all species had gene data on BLAST, particularly newly described species, some taxa were unable to be screened for contamination using BLAST. To identify potential contaminants in the samples that were difficult to BLAST, we determined sequence heterozygosity for all samples. To do this, we identified the longest sequence at each UCE locus using bioawk (https://github.com/lh3/bioawk?tab=readme-ov-file) and concatenated each of the longest sequences together to create a pseudo-genome for mapping purposes. Then, we used BWA-MEM (Li 2013) to index the constructed pseudo-genome and map the cleaned reads to the reference genome, followed by samtools (Danecek et al. 2021) to create a sorted BAM file. ANGSD (Korneliussen, Albrechtsen, and Nielsen 2014) was then used to calculate genome-wide heterozygosity for each sample. Samples with usually high heterozygosity (especially when compared with other samples of the same species or species group) were assumed to be contaminated and thus excluded from further analyses. One sample, *Pogonomyrmex marcusi* NK00, was kept despite high heterozygosity because *P. marcusi* is rarely collected and unlikely to be included in future molecular phylogenies of *Pogonomyrmex*.

### Data filtering, gene trees and loci selection

Contaminated samples as well as samples with 1,000 or fewer captured UCE loci were discarded, leaving 154 taxa. The command ‘phyluce_align_get_only_loci_with_min_taxa’ was used to select loci that were captured for 75% of the taxa, and produced 2,415 loci. To determine if these loci should be included in the concatenated tree inference, gene trees for each locus were inferred and assessed for relative bootstrap support. Loci were kept if the loci’s gene tree had over 80% bootstrap support of 80% of its internal nodes, resulting in a dataset of 2,304 loci. Subsequently, the 2,304 highly-supported loci were tested for symmetry using the IQ-TREE “—symtest-only” option; we were unable to significantly reject the symmetry of 1,423 loci, so these loci were included in the final dataset. These 1,423 loci were then concatenated into a PHYLIP file using ‘phyluce_align_concatenate_alignments’; the same 1,423 loci were used (in the form of individually-inferred gene trees) for the coalescent tree as well. The full 2,304 loci dataset of highly-supported gene trees was retained and used for maximum likelihood inference as well.

### Model selection, concatenated maximum likelihood tree inference, and coalescent summary tree inference

The best nucleotide substitution models were determined for each individual loci using ModelFinder (Kalyaanamoorthy et al. 2017); models used for each locus are available in the supplementary materials. Maximum likelihood analyses were performed using IQ-TREE 2.2.0 with 1,000 bootstrap replicates and each loci used as its own data partition (Minh et al. 2020). We used the 2304 high-supported gene trees to infer a coalescent summary tree using ASTRAL-III (Zhang et al. 2018).

### Divergence dating analysis

For the divergence dating analysis, the initial set of 154 taxa used in the IQ-TREE and ASTRAL inferences was pruned to 78 taxa. Outgroup species outside of Pogonomyrmecini were removed, and *Pogonomyrmex* species that were inferred to be monophyletic in the maximum likelihood phylogeny were pruned to one taxon per species. Species that were inferred to be non-monophyletic were excluded from the divergence time estimation.

To further reduce computational burden, we selected a small subset of loci from the larger 1,423-loci “good symmetry” dataset to use for the divergence date estimate. We used SortaDate (Smith, Brown, and Walker 2018) to determine the loci with the most ‘clocklike’ nucleotide evolution; the top 100 of these loci were selected to be used in divergence dating. The sequences of the top 100 loci were concatenated with Phyluce (“phyluce_align_concatenate_alignments”) and gene trees inferred for each of the 100 loci were coalesced into a species tree using ASTRAL-III, both of which were used as input data for divergence dating with MCMCTree (Yang 2007). We used the “approximate likelihood” method, a method that calculates the likelihood function during MCMC iteration and improves speed and efficiency of the MCMCTree run (Reis and Yang 2011). The tree root was calibrated using the range of ages in the 95% highest probability density for Pogonomrymecini (43.7-66.2 Mya) from the Myrmecine phylogeny (Ward et al. 2015).

### Historical biogeography and ancestral range reconstruction

To infer the historical biogeography of *Pogonomyrmex* and the Pogonomyrmecini tribe, we used the R (R Core Team 2021) package BioGeoBEARS (Matzke 2018) and implemented the best-fitting model determined by BioGeoBEARS, the Dispersal, Extinction, and Cladogenesis (DEC) model.We used seven discrete biogeographic regions defined by regionalization of the Neotropical and Neactic realms (Escalante, Rodriguez-Tapia, and Morrone 2021; Morrone et al. 2022) and the contemporary species range of each *Pogonomyrmex* species (Guénard et al. 2017; Janicki et al. 2016) included in the analysis. The biogeographic regions are indicated on a map in Figure 4 and include: 1) the entire Nearctic range of *Pogonomyrmex* and the Mexican transition zone (“Nearctic”), 2) lowland provinces of Mesoamerica (“Mesoamerica”), 3) the island of Hispaniola (“Hispaniola”), 4) the Pacific dominion excluding the Isthmus of Panama (“Pacific”), 5) the range of the cosmopolitan *Pogonomyrmex naegelii* which includes parts of the Boreal Brazilian Dominion, the South Brazilian Dominion, the Paraná Dominion, and the Brazilian Cerrado (“Brazilian”), 6) the Chacoan and Pampean provinces (“Chacoan”), and 7) the South American transition zone (“South American Transition Zone”).

## Results

### UCE Capture Statistics

Our final concatenated UCE sequence matrix, filtered to include samples with 1000 captured UCE loci or more, includes 154 samples representing 74 *Pogonomyrmex* species (∼80% of all described), 6 undescribed *Pogonomyrmex* species, outgroups from within the Pogonomyrmecini (*Hylomyrma* and *Patagonomyrmex* species), and four additional outgroup taxa (*Aphaenogaster occidentalis, Myrmica incompleta, Stenamma alas*, and *Veromessor lobognathus*). Out of the 2,524 loci that can be targeted for sequencing using the ant-specific hymenoptera probe set, we captured an overall mean of 2,232 UCE loci for the samples in the dataset (∼88% of loci). The final concatenated matrix consists of 2,304 UCE loci and is 1,693,317 base pairs; after filtering for symmetry, the final matrix consists of 1,423 UCE loci and is 1,039,348 base pairs long. While trees inferred with both the 2,304-loci and the 1,423-loci matrix have identical topologies, clades in the 1,423-loci matrix are more highly supported (Figure 2), and are shown in the main figures of the paper. The 2,304-loci tree is included in the supplement.

**Figure 2:**
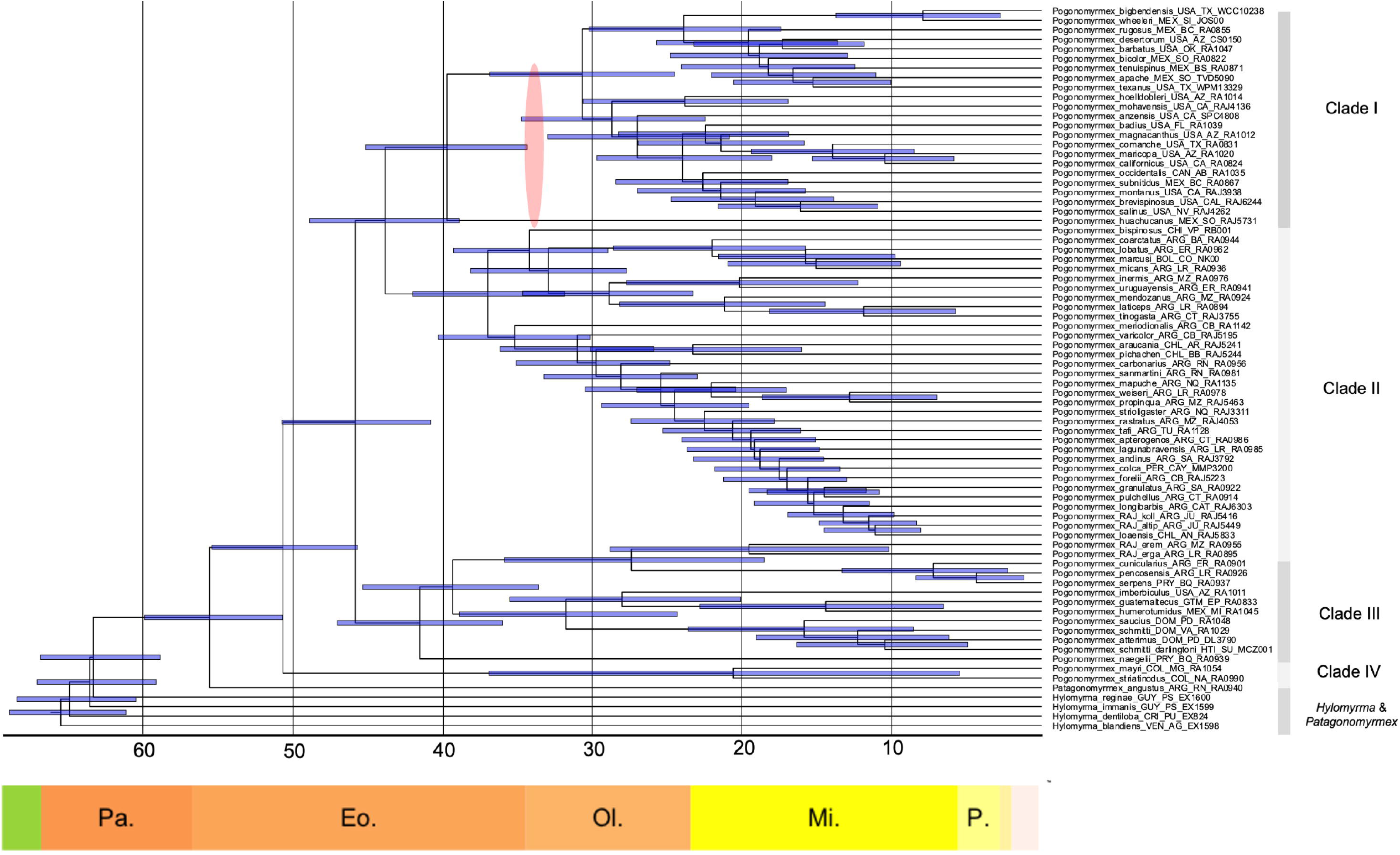
Maximum likelihood tree inferred using IQ-TREE 2 with the 1,423 “good symmetry” loci dataset. Background color indicates delimitation of the four main *Pogonomyrmex* clades inferred in the phylogeny, as well as Pogonomyrmecini outgroups *Hylomyrma* and *Patagonomyrmex*, labeled as such on the right side. *Pogonomyrmex* species groups are indicated to the right of the gray lines. Taxa names colored in red indicate taxa formerly placed in the genus *Ephebomyrmex*; one taxa name, that of *Pogonomyrmex mayri*, is colored blue to indicate its former placement in the genus *Forelomyrmex*. Nodes without support values all received maximum bootstrap support while blue circles indicate nodes with bootstrap support <95% and corresponding bootstrap values are displayed to the left of each circle.

### Phylogenomic inference and evolutionary relationships

The concatenated maximum likelihood tree inferred with IQ-TREE 2 and the coalescent species tree inferred using ASTRAL-III all strongly support four monophyletic *Pogonomyrmex* clades. These clades include, and will be referred to in this paper as: Clade I, which contains 26 species exclusively found in North America; Clade II, which contains 36 species exclusively found in South America; Clade III, which consists of 12 species from North America, Mesoamerica, Hispaniola, and South America, many of which are species formerly classified as the genus *Ephebomyrmex*; and Clade IV, a small clade that includes only *P. mayri* and *P. striatinodus* and diverged earlier from the rest of *Pogonomyrmex*.

*Pogonomyrmex* species groups, determined by morphology and described in multiple publications (Cole 1968; Johnson 2015, 2021; Johnson and Cover 2015; Johnson, Overson, and Moreau 2013; MacKay 1981; Vásquez-Bolaños and Mackay 2004) were overall supported by both inferred tree topologies. In particular, the monophyly of the *P. rastratus*-group, a large species group containing many newly described species and some yet-to-be-described species, was supported by all tree topologies. However, two current *Pogonomyrmex* species groups, the *Pogonomyrmex bispinosus*-group and the *Pogonomyrmex californicus*-group were not supported by our UCE tree topologies.

The majority of species with two or more samples included in the two inferred trees are recovered as monophyletic. Still, a number of species were not inferred to be monophyletic; most of these species are within the *Pogonomyrmex rastratus*-group clade. Branches within the *P. rastratus*-group clade are among the shortest and have the poorest bootstrap and posterior probability support values of the entire tree. Outside of the *P. rastratus*-group clade, the tree topology is very well supported, with an average bootstrap support value of 99.2% (96.2% including the *P. rastratus*-group clade).

### Divergence dating

Our MCMCTree divergence dating analysis inferred a Paleocene origin for Pogonomyrmecini with a mean crown age of 65.2 million years old (Figure 3). This is the oldest the group has been inferred in all divergence dating analyses including the tribe; Ward et al. (2015b) inferred the crown age of Pogonomyrmecini to be 54.3 million years old, and a very recent UCE phylogeny of ants inferred a ∼45 million year old origin Pogonomyrmecini (Borowiec et al. 2024)) The mean age of *Pogonomyrmex* is inferred to be 50.4 million years old, placing the origins of the group in the early Eocene. The mean ages of the four large clades, Clade IV, Clade III, Clade II, and Clade I, were inferred to be 50.4 Mya, 41.3 Mya, 39.9 Mya, and 37.0 Mya, respectively.

**Figure 3:**
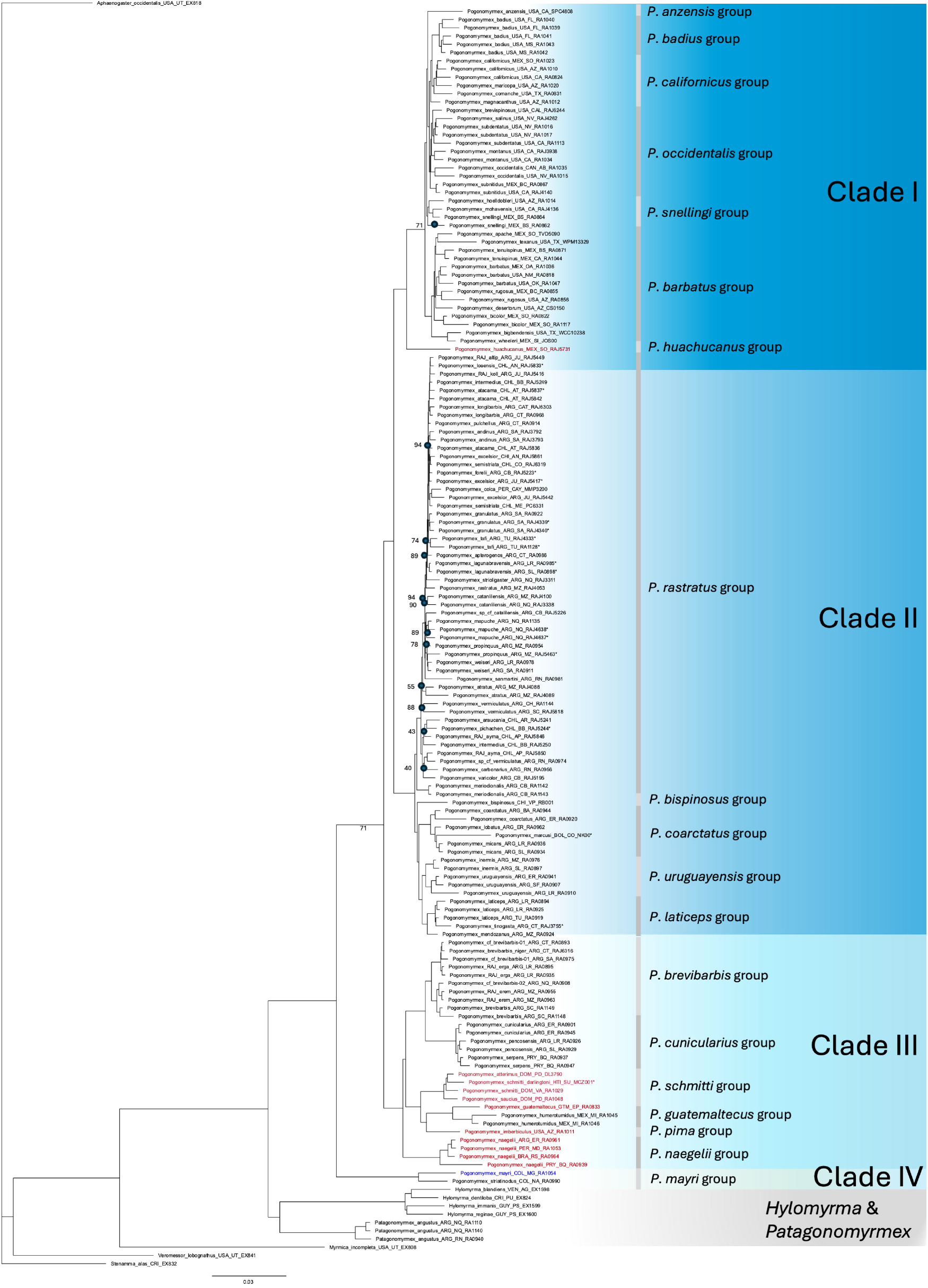
Chronogram showing the divergence dating analysis performed with MCMCTree. Absolute time in millions of years is indicated on the x-axis, and geologic epochs are indicated underneath the absolute time. Node bars indicate the 95% HPD range for the age of each node. The translucent red oval indicates the approximate age of the sole *Pogonomyrmex* fossil, *Pogonomyrmex fossilis*, as well as the fossil’s possible placements on branches in North American *Pogonomyrmex* clade.

### Historical biogeography

The ancestral range reconstruction performed on the dated phylogeny with BioGeoBEARS infers a “Pacific dominion” origin for both Pogonomyrmecini and *Pogonomyrmex* (Figure 4). However, the ancestors of all *Pogonomyrmex* species excluding the *Pogonomyrmex mayri* clade are inferred to have originated in the “South American Transition Zone” biogeographic region. Clade I, the North American clade, is inferred to have dispersed to North America in a single event in the late Eocene, and a group within the Clade III is inferred to have dispersed to Hispaniola, Mesoamerica, and North America within the Oligocene and Miocene. The most recent dispersals, dispersals to the “Chacoan” biogeographic region, were inferred to have occurred during the Miocene and the Pliocene.

**Figure 4:**
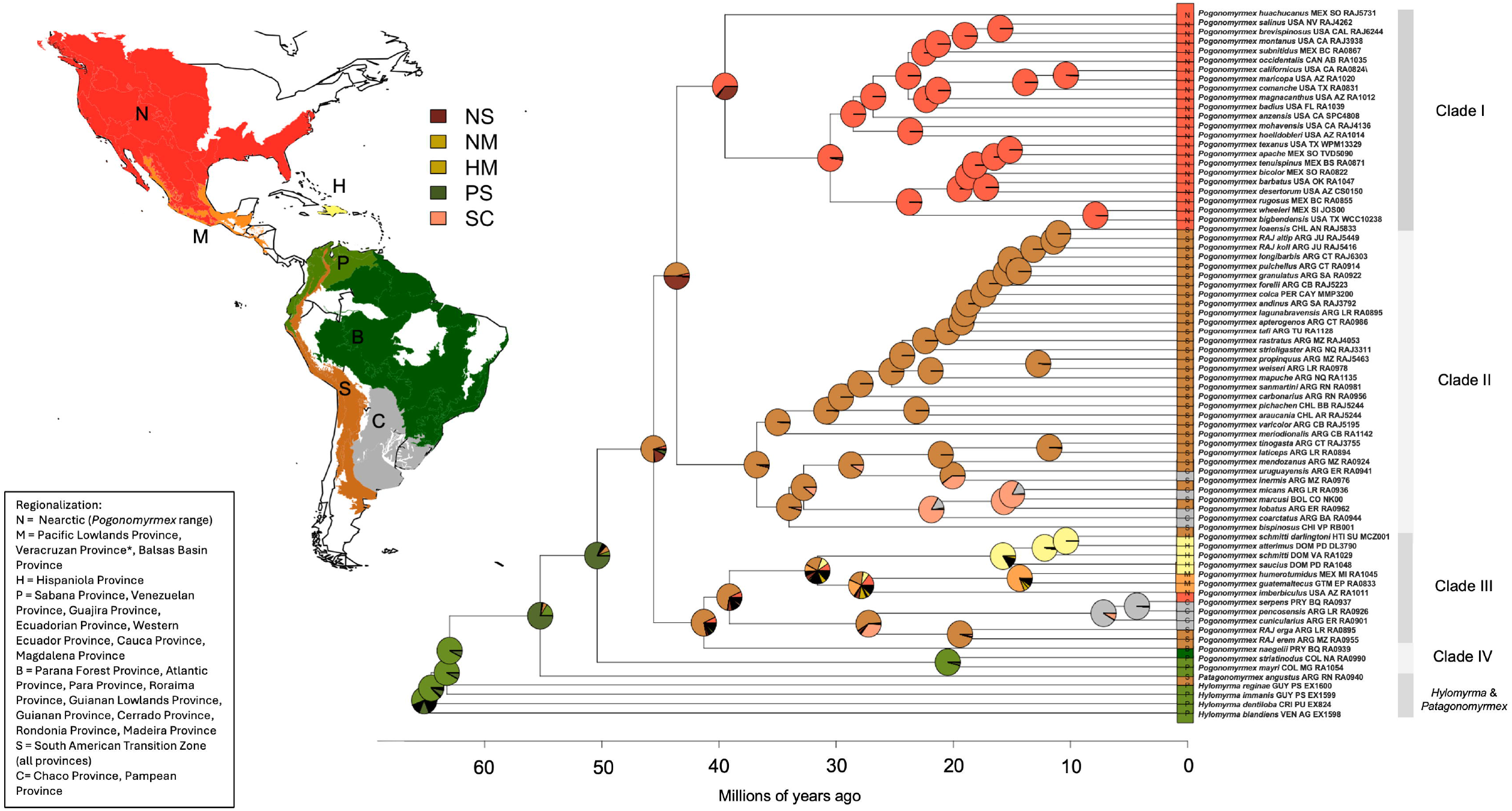
Ancestral range reconstruction from BioGeoBEARS using dated MCMCTree chronogram. The four major *Pogonomyrmex* clades are delimited by shaded columns to the right of the tree. Map on the left indicates the seven discrete ranges assigned to each of the tips, labeled with colors and letters (N=“Nearctic”, M=“Mesoamerica”, H=“Hispaniola”, P= “Pacific”, B=“Brazilian”, C=“Chacoan”, S=“South American Transition Zone”). Colored boxes to the right of the map indicate intermediate ranges constructed by BioGeoBEARS. Regionalization, consisting of combined provinces defined by Morrone et al. (Morrone et al. 2022), is indicated in the inset in the bottom left corner.

## Discussion

This study is the first to utilize phylogenomic data to infer the evolutionary relationships of species within the harvester ant genus *Pogonomyrmex*; our inferred trees are the most comprehensive to date and include the most *Pogonomyrmex* species of any phylogenetic inference. Though our inferred tree topologies are generally consistent with the existing morphological and molecular phylogenies of the genus (Johnson and Moreau 2016; Parker and Rissing 2002; Taber 1999), our increased taxon and genomic sampling has revealed new relationships within *Pogonomyrmex* that contradict some species group classifications based on morphology. Further, we inferred an earlier origin of *Pogonomyrmex* and the Pogonomyrmecini tribe than previous work, as well as a northern South American origin for both groups.

Prior to this work, the *Pogonomyrmex* phylogeny has been inferred using morphological characters (Taber 1999), a single mitochondrial gene (Parker and Rissing 2002), and six concatenated genes (one mitochondrial gene and five nuclear genes) (Johnson and Moreau 2016). Previous phylogenies have only included a portion of described *Pogonomyrmex* species, with the two earliest phylogenies representing mainly Nearctic taxa. Johnson and Moreau’s (2016) phylogeny included substantially more sampling from South America and Hispaniola; the four major clades inferred in our UCE phylogeny were also inferred in the six-gene phylogeny. Further, most of the species-level relationships within these major clades are also inferred as being similar between the six-gene and the UCE phylogeny. *Pogonomyrmex huachucanus*, a former *Ephebomyrmex* species, is inferred as being sister to the rest of the North American *Pogonomyrmex* clade, a result we corroborate as seen by Johnson and Moreau’s (2016) phylogeny as well as Parker and Rissing’s 2002 mitochondrial gene phylogeny.

However, the inclusion of many more *Pogonomyrmex* species in our UCE phylogeny compared to previous phylogenies (74 species in the UCE phylogeny; 28 species in Johnson and Moreau 2016) has called into question some species group relationships previously determined by morphology and ecological similarity. The *P. bispinosus*-group is one such example; *P. bispinosus* was inferred as sister to a large clade of South American *Pogonomyrmex* that includes the *P. coarctatus*-group, the *P. laticeps*-group, and *P. bispinosus*-group species *P. inermis* and *P. uruguayensis*. Since no other phylogenetic inference has included all three *P. bispinosus-*group species, these species group relationships should be re-examined considering new molecular evidence. The *P. californicus*-group is also not supported by the tree topologies. Three *P. californicus*-group species, *P. hoelldobleri, P. mohavensis*, and *P. snellingi*, are inferred to form a clade that is sister to the rest of the *P. californicus*-group, the *P. occidentalis*- group, and *P. badius. Pogonomyrmex anzensis* is inferred as sister to these species groups as well. The other four of the *P. californicus*-group’s eight species, *P. magnacanthus, P. comanche, P. californicus*, and *P. maricopa*, are inferred to share a common ancestor, and are the closest sister group to the *P. occidentalis*-group. This *P. californicus*-group was also inferred to be polyphyletic in Johnson and Moreau’s (2016) molecular phylogeny, though the relationships of the *P. californicus*-group species to those in the *P. barbatus*-group differs in our phylogeny.

Our ancestral range reconstruction infers both Pogonomyrmecini and *Pogonomyrmex* to have origins in northern South America. This finding is consistent with earlier theories about the geographic origins of *Pogonomyrmex* based on comparative morphology, species distribution, and geologic trends (Fernández and Palacio 1997; Kusnezov 1951). The ancestors of *Pogonomyrmex* probably lived in humid environments and were likely not granivorous or very infrequently granivorous, much like species in Clade IV (the *Pogonomyrmex mayri-*group) today. Climatic cycles, known to cause cyclical vegetational changes in South America for the last 60 million years, may have caused these humid areas to become cooler and drier for stretches of time (Haffer 1996). It is in these newly open areas that the ancestors of all other *Pogonomyrmex* besides the *P. mayri* clade may have adapted to relying on seeds as a primary food source (Fernández and Palacio 1997). These xeric-adapted, granivorous *Pogonomyrmex* ancestors were then able to spread within South America, while the ancestors of the *Pogonomyrmex mayri*-clade (*Pogonomyrmex mayri, Pogonomyrmex striatinodus*, and likely *Pogonomyrmex sylvestris* and *Pogonomyrmex stefani*, taxa not included in our phylogeny) were confined to forest refugia, as the *P. mayri* clade still is today.

During the middle-to-late Eocene, *Pogonomyrmex* dispersed to the Nearctic region, giving rise to the Clade I, the last common ancestor of which lived about 40 million years ago. The ancestors of the Nearctic *Pogonomyrmex* may have spread to North America via the Panama Arc, a chain of ephemeral volcanic islands formed by the subduction of the Pacific-Farallon Plate beneath the Caribbean and South American plates that would later move eastward to form the Isthmus of Panama (O’Dea et al. 2016). During the late Eocene, the Panama Arc was nearly in its current position, but the South American plate had not yet collided with the Caribbean plate, so dispersal by flight would have been necessary to cross the island arc. Much of North American *Pogonomyrmex* diversification likely occurred in the Oligocene and Miocene; the Oligocene climate of North America was cooler and drier than compared to the Eocene, particularly in the region where the only known fossil of the genus, *Pogonomyrmex fossilis*, was discovered in present-day Colorado, USA. Grasses, a prominent food source for *Pogonomyrmex*, were likely already diverse in arid environments of Oligocene North America (Pimentel et al. 2017). The further aridification of southwest and south-central North America, caused by the uplift of the Rocky Mountains, likely selected for the desert-adapted *Pogonomyrmex* species known in those regions today (Blakey et al. 2018).

Clade III split from a common ancestor with Clades I & II in the middle Eocene. Some members of this clade, the progenitors of the *Pogonomyrmex pima*-species group, the *Pogonomyrmex schmitti*-group, and the *Pogonomyrmex guatemaltecus*-species group, left South America and likely dispersed to Central America via the Panama Arc sometime in the middle Oligocene. Members of the *Pogonomyrmex pima*-species group soon migrated to North America. Later, by the middle Miocene, the ancestors of the *Pogonomyrmex schmitti*-group dispersed to Hispaniola. Dispersal from Central America to the Greater Antilles has been inferred in other groups, including *Platythyrea* ants that are inferred to have also dispersed to the Greater Antilles by the middle Miocene (Crews and Esposito 2020), as well as two groups of beetles (Browne, Peck, and Ivie 1993; Liebherr 2019). The *Pogonomyrmex schmitti*-group likely dispersed to Hispaniola via Cuba, as this route provides the shortest distances to travel over water; however, no *Pogonomyrmex* species are presently found on Cuba. The respective presence and absence of *Pogonomyrmex* in Hispaniola and Cuba may be consistent with a pattern of observed species diversity in some taxa being higher in Hispaniola than in Cuba, a trend hypothesized to be caused by differences in topological complexity between the two islands (Crews and Esposito 2020).

The *Pogonomyrmex rastratus*-group clade, the largest species group clade in our phylogeny, contains the most *Pogonomyrmex* species that are inferred to be younger than 20 million years old. The clade also has some of the most poorly supported species-level relationships of the tree, and of the fourteen *P. rastratus*-group species that were represented by two or more samples, seven of these species were not inferred to be monophyletic. While this ambiguous topology could potentially be caused by undetected contamination or sequence degradation, it is also possible that lineages within the *P. rastratus-*group have incompletely sorted. Speciation within this group may be driven by new niche formation caused by the ongoing Andean uplift; species within this group are often geographically close to one another but are separated by elevational differences. Introgression and even more recent hybridization between species in this group are both possible explanations for the lack of clarity in species-level relationships; North American species *Pogonomyrmex barbatus* and *Pogonomyrmex rugosus*, species that we have inferred to have last shared a common ancestor 20 million years ago, often hybridize, which has been studied extensively (Cahan, Nguyen, and Zhou 2022; Mott, Gadau, and Anderson 2015). Population genetics with increased sampling of multiple individuals of *P. rastratus*-group species may be necessary to understand the evolutionary relationships within this species group.

While some *Pogonomyrmex* species are found outside of deserts, the greatest species diversity of the genus in the North and South American deserts resemble other desert amphitropical disjunctions observed in both plants (Lia et al. 2001; Wen and Ickert-Bond 2009) and animals (Oberski 2022; Wilson, Carril, and Sipes 2014). Like *Pogonomyrmex, Dorymyrmex* ants (Oberski 2022), *Diadasia* bees (Wilson et al. 2014), and *Larrea* creosote bush (Lia et al. 2001) are all inferred to have originated in South America, to have dispersed to North American deserts, and to have subsequently diversified. However, both *Dorymyrmex* and *Diadasia* are inferred to have dispersed to North America during the Miocene—*Diadasia* also putatively crossed via the pre-isthmus Panama Arc—but the presence of a fossil from the genus suggests that *Pogonomyrmex* was certainly on the North American continent by the Oligocene, and our divergence dating analysis suggests an even earlier dispersal. Unlike other amphitropically disjunct desert species, *Pogonomyrmex* did not move from one desert to another; instead, it is more likely that *Pogonomyrmex* species on both continents convergently evolved to succeed in both deserts, having had traits that made them already competent competitors in arid environments.

Overall, our results suggest that adaptation to arid environments was a key innovation for *Pogonomyrmex*, allowing them to spread and diversify within the Americas while the climate was cooling and arid grasslands and deserts were actively being formed. These innovations could include a transition to granivory and the development of the psammophore for digging nests in sandy soils, both traits present in nearly all *Pogonomyrmex* species, but most pronounced in *Pogonomyrmex* species living in the most arid environments. Studying the evolution of these adaptations to arid environments will be necessary to understand more about the evolution and diversification of this prominent desert ant.

## Acknowledgments

We thank Chloe Jelley for comments on earlier versions of this manuscript which greatly improved it. L.C.G. thanks Dr. Miles Zhang and Dr. Jacob Landis for help with contaminant identification.

## Funding

This material is based upon work supported by the National Science Foundation Graduate Research Fellowship (NSF DGE 2139899 to L.C.G.). This work was also supported by the National Science Foundation (NSF DBI 2210800 to C.S.M.).

## Authorship Contributions

L.C.G.: Conceptualization; Data curation; Formal analysis; Investigation; Methodology; Project administration; Software; Validation; Visualization; Writing—original draft. R.A.J.: Conceptualization; Data curation; Resources; Writing—review & editing. C.S.M.: Conceptualization, Funding acquisition, Resources, Supervision, Writing—review & editing.

## Data Accessibility

Raw sequence reads in FASTQ format are deposited in the NCBI Sequence Read Archive (accession #PRJNA1177438).

## Notes

### Competing Interest Statement

The authors have declared no competing interest.

